# Beyond a single threshold: calibration of CADD v1.7 and AlphaGenome for ACMG/AMP computational evidence across variant classes, model versions, and reference data

**DOI:** 10.64898/2026.10.31.763309

**Authors:** José Patricio Miranda, Lorena Carvajal, Ignacio Aldana, Pablo Alarcón, Cristian Saavedra

## Abstract

**Purpose:** Computational predictors provide PP3 and BP4 evidence under the ACMG/AMP guidelines, but that evidence holds only within the conditions of its calibration. We asked how far a calibrated threshold travels.

**Methods:** Following a plan fixed before any result, we used the ClinGen local posterior probability method to calibrate CADD v1.7 (Combined Annotation Dependent Depletion) and the AlphaGenome Variant Impact score (AVI) on rare ClinVar missense variants, then changed one condition at a time: variant class, model version, reference set, and gene.

**Results:** Both predictors reached moderate pathogenic and strong benign evidence, and their thresholds held for variants classified after calibration. AVI outperformed CADD in every comparison. Thresholds available in 2024 gave AVI evidence in the final direction for 72.5% of the variants of uncertain significance resolved by 2026, against 60.5% for CADD. Missense thresholds gave false benign evidence to pathogenic noncoding variants. Moving from CADD v1.6 to v1.7 altered the evidence level of a third of variants without improving discrimination, and changing the reference set moved the thresholds.

**Conclusion:** A calibrated threshold belongs to a predictor, a variant class, a model version, and a reference set together. Laboratories should report the source of every threshold they apply.

## INTRODUCTION

The ACMG/AMP guidelines classify germline variants into five categories by weighing evidence from population data, computational predictions, functional studies, segregation, and de novo occurrence.^1^ Computational evidence supports PP3 when predictors indicate a damaging effect and BP4 when they suggest the variant is unlikely to be harmful. Both criteria are limited to supporting strength, and the guidelines leave laboratories to choose their own tools and thresholds. As a result, approaches vary between laboratories.^2^ This inconsistency can affect how variants of uncertain significance (VUS) are interpreted. In a study of 1.7 million people tested for hereditary disease, 41% had at least one VUS. Most were missense variants, and VUS were more common among people of non-European ancestry.^3^

The Bayesian formulation of the ACMG/AMP framework gives each evidence strength an odds of pathogenicity, against which any quantitative predictor can be calibrated.^4,5^ The ClinGen Sequence Variant Interpretation working group built on it a local posterior probability method that uses ClinVar pathogenic and benign variants as labels, rare gnomAD variants to size the estimation windows, and a prior probability of pathogenicity for rare missense variants.^6^ REVEL, BayesDel, and VEST4 reached PP3 strong and BP4 strong or very strong, and clinical laboratories and ClinGen expert panels adopted the resulting intervals, which later work extended to AlphaMissense.^6–8^

In routine practice, genome-wide predictors are used when missense-specific tools cannot be applied, including splice-region, intronic, untranslated, and synonymous variants. Laboratories and their interpretation software often rely on CADD for these variants, frequently using a single cutoff for each variant type.^9^ Released in 2024, CADD v1.7 incorporates features from protein language models and regulatory convolutional networks.^10^ Because its PHRED score is a genome-wide ranking, the v1.6 thresholds (PP3 ≥25.3 for supporting evidence and ≥28.1 for moderate evidence)^6^ should not be assumed to apply to v1.7. In 2026, AlphaGenome added sequence-based predictions of regulatory, splicing, and protein-level effects,^11^ which were combined with protein and conservation features into the AlphaGenome Variant Impact score (AVI) (Cheng J, Taylor KR, Nicolaisen L, et al, medRxiv preprint, 2026, doi:10.64898/2026.09.16.26363192).

Three features of current reference data can bias a calibration. Predictors look better than they are when tested on variants whose classifications relied on them,^12^ and calibrated predictors have been cited as PP3 or BP4 evidence in ClinVar submissions since 2022. The original calibration removed variants listed in the Human Gene Mutation Database (HGMD),^13^ which most laboratories cannot repeat because the professional database requires a license. Finally, a genome-wide threshold assumes a uniform relationship between score and pathogenicity across all genes, even though genes differ in constraint, inheritance, and disease mechanisms.^14,15^ Multiplexed assays of variant effect (MAVEs) measure the functional impact of thousands of variants per gene without relying on clinical classifications.^16^ They have been used to benchmark predictors^17^ and as functional evidence,^18,19^ but not to test PP3/BP4 thresholds. We calibrated CADD v1.7 and AVI using an implementation that reproduces the published CADD v1.6 thresholds, under an analysis plan fixed before any results were available. We then tested whether the thresholds hold for variants classified after calibration and would have given correct evidence for VUS that were later resolved, identified the current VUS that receive enough concordant evidence to approach a new classification, evaluated whether missense thresholds can be applied to other variant types, and measured how much the thresholds depend on the model version, the reference data, and the gene.

## MATERIALS AND METHODS

### Study design

The reference sets, calibration parameters, sensitivity analyses, and discrimination metrics were defined before any result from CADD v1.7 or AVI was obtained, and the plan was committed to the project’s version-controlled repository on September 25, 2026. Each later analysis was registered as a dated amendment before it was run (Table S1). Three amendments followed results already observed, namely the paired comparison of AVI and CADD, the analysis of threshold drift across ClinVar releases, and the HGMD exclusion.

### Predictors and reference sets

CADD v1.7 GRCh38 PHRED scores for all possible SNVs were downloaded from the official website,^10^ and AVI Phred scores were obtained from the AlphaGenome Atlas (Cheng J et al, medRxiv preprint, 2026). The comparators were REVEL v1.3,^20^ BayesDel without allele frequency,^21^ and AlphaMissense.^22^ All possible SNVs in the coding sequences of the 19,229 GENCODE v50 MANE Select transcripts were enumerated and annotated.^23,24^ As in the original calibration,^6^ the labeled set comprised ClinVar GRCh38 SNVs (release of September 23, 2026)^25^ that are missense in the MANE Select transcript, classified as Pathogenic, Likely pathogenic, or both (P/LP) or as Benign, Likely benign, or both (B/LB) with at least one review star, with a gnomAD v4.1^26^ allele frequency below 0.01, and located in a disease gene (at least one P/LP record with at least one star). The unlabeled set, used only to size the estimation windows, comprised rare MANE missense SNVs present in gnomAD v4.1 and absent from ClinVar in the same genes, with frequencies from the Ensembl VEP release 116 cache.^27^ The sets contained 59,979 P/LP and 77,128 B/LB variants across 4,826 genes, as well as 1,955,995 gnomAD variants. HGMD-listed variants were retained because their effects were tested separately, and training variants were retained because neither CADD nor AVI is trained on ClinVar labels. Individual variants are described on the MANE Select transcript with its RefSeq accession and version and on GRCh38 (NC_000019.10 for *LDLR*, NC_000001.11 for *PIK3CD*), and every variant description reported here was verified with VariantValidator.^28^

### Calibration

We ported the original MATLAB implementation of the local posterior probability method^6^ to R, keeping its parameters: an estimation window that grows in steps of the predictor’s score increment (Table S2) until it holds at least 100 labeled and 3% of unlabeled variants, a prior of 0.0441, and the likelihood ratios 2.41, 5.79, 33.5, and 1,124 for supporting to very strong PP3 with their reciprocals for BP4.^4–6^ Each threshold is the most extreme score of the continuous region meeting the required ratio, taken as the one-sided 95% bound from 10,000 bootstrap replicates, and a level was considered not reached if more than 5% of replicates failed to meet it. On the original authors’ deposited data, the code reproduced the published CADD v1.6 thresholds and showed that thresholds in sparse score regions depend on the random seed (Figures S1 and S2, Tables S3 and S4), so each main threshold is the median over five prespecified seeds. The four sensitivity analyses are described in the Supplemental Methods (Table S5). Discrimination was measured by the area under the receiver operating characteristic curve (AUROC) with 1,000 bootstrap replicates and paired bootstrap tests.

### Temporal hold-out and resolved VUS

Both predictors were recalibrated on reference sets built in the same way from the ClinVar release of January 27, 2024, the last one a laboratory could have used before CADD v1.7 was released. For each evidence interval, we compared the likelihood ratio among the 22,626 P/LP and 36,294 B/LB variants of the 2026 sets that were absent in 2024 with the ratio its strength requires. We then followed every rare missense VUS with at least one star in a disease gene from the January 2024 release through the September 2026 release, and considered it resolved if it was classified as P/LP or B/LB. Evidence from all five predictors was assigned using thresholds calibrated only to the 2024 release, as a laboratory could have applied them at the time. For each predictor, we report the proportion of resolved variants with evidence in the final direction, the ACMG/AMP points contributed (±1 supporting, ±2 moderate, ±4 strong),^5^ and the likelihood ratio of each interval. Because 2026 classifications may cite REVEL, BayesDel, AlphaMissense, or CADD, we used AVI, which no submitter could have used, as the negative control for circularity.

### Current VUS and version transfer

We assigned evidence to the 1,208,461 missense VUS with at least one star in disease genes of the September 2026 release, using the thresholds of Table 1 for CADD v1.7 and AVI and the published ClinGen thresholds for REVEL, BayesDel, and AlphaMissense.^6,8^ Evidence was considered concordant when the three missense predictors and AVI pointed in the same direction, and took the smallest absolute points among the three missense predictors, the conservative choice for a laboratory applying one predictor. Using the ACMG/AMP point system,^5^ we report, for each VUS, the remaining evidence needed to complete a likely pathogenic or likely benign classification. ClinVar does not record those points, so this is a scenario rather than a reclassification. To measure the effect of transferring thresholds between model versions, we also read the CADD v1.7 scores with the published CADD v1.6 thresholds. Counts refer to the 1,208,430 VUS with allele frequency below 0.01, and no variant was removed from the lists.

**Table 1.** Calibrated PP3/BP4 thresholds for CADD v1.7 and AVI (rare missense variants, ClinVar 2026).

| Evidence | Required LR | CADD v1.7 PHRED | CADD P/LP % | CADD B/LB % | AVI Phred | AVI P/LP % | AVI B/LB % | CADD v1.6 PHRED (Pejaver et al. 2022) | CADD v1.6 PHRED (2026 data) |
|---|---|---|---|---|---|---|---|---|---|
| PP3 Very strong | 1124 | not reached | — | — | not reached | — | — | not reached | not reached |
| PP3 Strong | 33.53 | not reached | — | — | not reached (reached in 59% of replicates) | — | — | not reached | not reached |
| PP3 Moderate | 5.79 | ≥ 28.0 | 46.2 | 4.8 | ≥ 26.44 | 67.9 | 4.0 | ≥ 28.1 | ≥ 27.5 |
| PP3 Supporting | 2.406 | ≥ 25.6 | 26.1 | 6.6 | ≥ 25.18 | 12.1 | 3.1 | ≥ 25.3 | ≥ 25.2 |
| BP4 Supporting | 1/2.406 | ≤ 22.6 [22.6–22.7] | 4.1 | 16.1 | ≤ 22.95 | 4.3 | 18.7 | ≤ 22.7 | ≤ 22.9 |
| BP4 Moderate | 1/5.79 | ≤ 19.59 | 2.1 | 29.1 | ≤ 20.52 [20.52–20.53] | 2.7 | 38.8 | ≤ 17.3 | ≤ 21.1 |
| BP4 Strong | 1/33.53 | ≤ 8.34 | 0.3 | 23.8 | ≤ 9.15 [9.15–9.17] | 0.3 | 22.4 | ≤ 0.15 | ≤ 7.35 |
| BP4 Very strong | 1/1124 | not reached | — | — | not reached | — | — | not reached | not reached |

Thresholds are one-sided 95% bootstrap bounds (10,000 replicates) of the ClinGen local posterior probability method (prior 0.0441), median [range] over five seeds. PP3 applies at or above the threshold; BP4 applies at or below the threshold. The reference set comprised 59,979 P/LP and 77,128 B/LB rare MANE Select missense ClinVar variants (4,826 genes) and 1,955,995 rare gnomAD v4.1 missense variants. P/LP % and B/LB % give the percentage of reference variants at each level (in-sample). LR, likelihood ratio required for each strength. AVI PP3 strong was reached in 59% of bootstrap replicates (95% required). CADD v1.6 thresholds from Pejaver et al.<sup>6</sup>, reproduced from the original data with our implementation (Table S3). The last column gives CADD v1.6 recalibrated on the same 2026 reference sets (median of five seeds).

### Comparators, ClinVar releases, and HGMD exclusion

The five predictors were calibrated on the reference variants scored by all of them (59,034 P/LP, 75,306 B/LB, and 1,888,558 gnomAD variants). To examine drift, we built reference sets from the ClinVar releases of December 2019 (used in the original study), November 2022, January 2024, and September 2026, with AVI as the negative control. Thresholds were compared across scales as the percentage of gnomAD variants meeting them. For the HGMD analysis, we removed from the 2019 and 2026 sets every SNV at a position that matches an HGMD Public (release 2020.4) record in the Ensembl VEP cache, and recalibrated all predictors.

### Functional assays, gene classes, and variant classes

We gathered all public human MaveDB^16^ score sets whose calibration separates functionally normal from abnormal variants and computed, for each calibrated interval, the ratio P(interval | abnormal) / P(interval | normal) and the AUROC. An abnormal assay result does not imply pathogenicity, so this analysis tests the direction and order of the intervals, not the ClinGen ratios. Genes were grouped by constraint (gnomAD v4.1 LOEUF deciles 0–2, 3–6, and 7–9),^14^ mode of inheritance (ClinGen gene–disease validity^29^ or the Gene Curation Coalition^30^), and disease mechanism (Gene2Phenotype^31^), and each class was calibrated with the global prior. ClinVar variants outside the MANE coding sequence were grouped into region classes; the missense thresholds were evaluated within each class and on saturation mutagenesis MPRA data for 20 regulatory elements,^32^ and the intronic splice region (3–8 nucleotides from an exon) was calibrated separately. To separate the model version from the reference data, CADD v1.6 and v1.7 were calibrated on the same 2,093,102 reference SNVs.

### Software

Analyses were performed in R 4.4.2 with Bioconductor 3.20, data.table, ggplot2, and openxlsx, with package versions fixed in a renv lockfile.

## RESULTS

### Calibrated thresholds for CADD v1.7 and AVI

With 59,979 P/LP and 77,128 B/LB variants from 4,826 genes, CADD v1.7 reached PP3 supporting at PHRED ≥25.6 and PP3 moderate at ≥28.0, and BP4 supporting, moderate, and strong at ≤22.6, ≤19.59, and ≤8.34 (Table 1, Figure 1). AVI reached PP3 supporting and moderate at Phred ≥25.18 and ≥26.44, and BP4 supporting, moderate, and strong at ≤22.95, ≤20.52, and ≤9.15. Neither predictor reached PP3 strong. The AVI posterior leveled off just below the required value, crossing it in 59% of bootstrap replicates, and AVI reached PP3 strong (≥30.23) when the reference set was restricted to variants with two or more stars.

**Figure 1.**
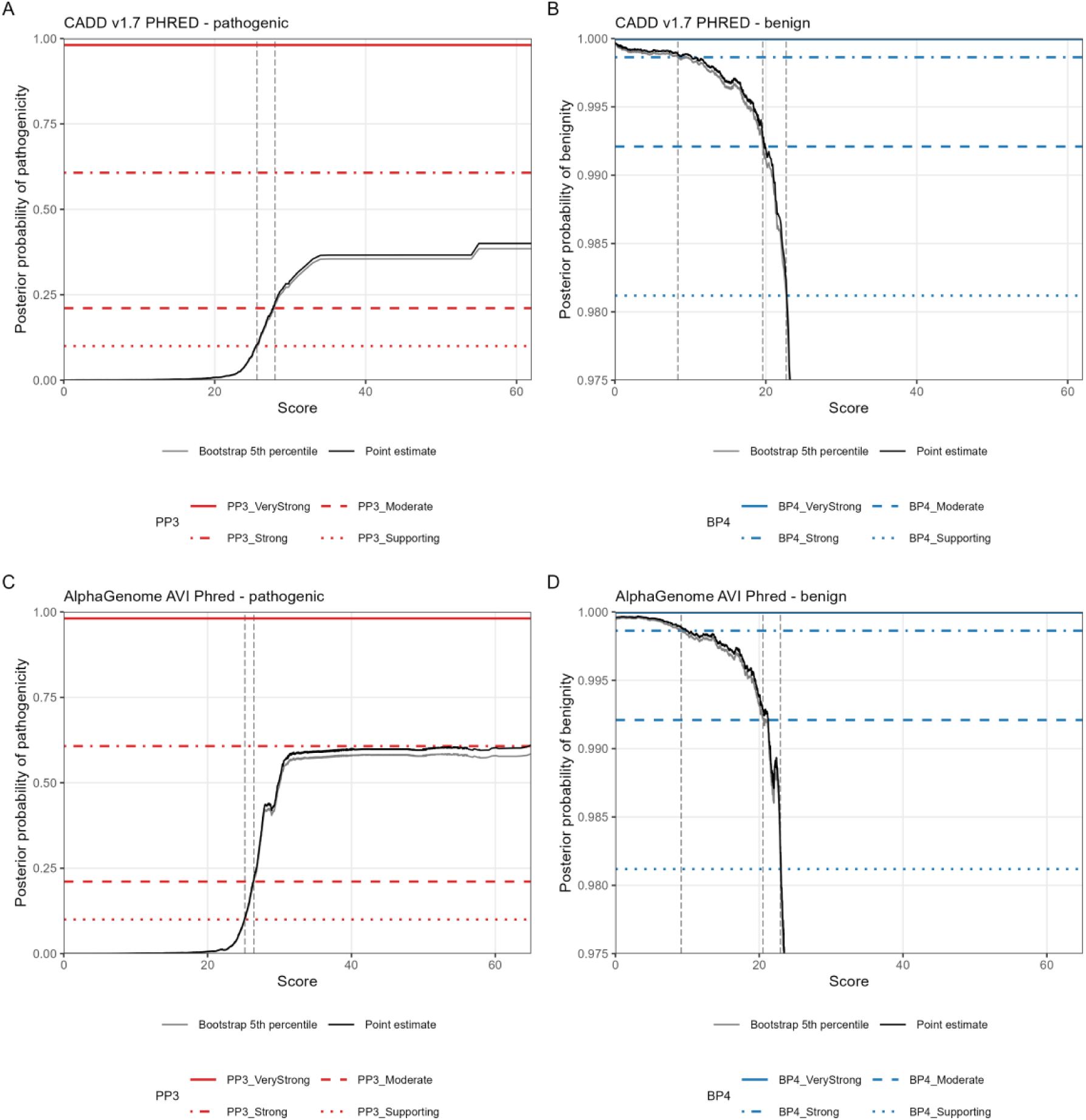
Local posterior probability of pathogenicity and benignity for CADD v1.7 and AVI. (A and B) CADD v1.7 PHRED. (C and D) AlphaGenome Variant Impact (AVI) Phred. Panels A and C show the posterior probability of pathogenicity, and panels B and D show the posterior probability of benignity as a function of the score, estimated with the ClinGen local posterior probability method on rare MANE Select missense variants from ClinVar (release of September 23, 2026) and gnomAD v4.1. Black lines are the point estimates, and grey lines are the 5th percentile of 10,000 bootstrap replicates (seed 20260925). Horizontal lines mark the posterior probability required for each evidence strength (supporting, moderate, strong, and very strong), and dashed vertical lines mark the final thresholds, defined as the one-sided 95% bootstrap bound.

Thresholds calibrated on the January 2024 release differed from the 2026 thresholds by 0.6 units or less, and every evidence interval of both predictors met the required likelihood ratio among the 58,920 variants classified after January 2024 (Figure S3, Table S6).

### Comparison with missense-specific predictors

AVI separated P/LP from B/LB variants better than CADD v1.7 (AUROC 0.940 versus 0.902). On the variants scored by all five predictors, REVEL, BayesDel, and AlphaMissense discriminated better still (AUROC 0.949–0.961) and were the only predictors to reach PP3 strong (Figure S4, Table S7). On functional data, this advantage disappeared. In 93,154 MANE missense SNVs from 39 genes, with functional classes from 89 MaveDB score sets (Table S8), the intervals for each predictor pointed in the expected direction. They followed the order of evidence strength (Table S9). AVI (AUROC 0.764) discriminated as well as BayesDel (0.766) and REVEL (0.757), behind AlphaMissense (0.782) and ahead of CADD (0.725), which it exceeded in 32 of the 36 genes with enough variants (Figure S5).

### VUS resolved after calibration

Of 657,184 rare missense VUS in disease genes in the January 2024 release, 1,790 (0.3%) had been classified as P/LP and 3,989 (0.6%) as B/LB by September 2026 (Table S10). With thresholds calibrated on the 2024 release alone, AVI gave evidence in the final direction for 72.5% of these variants and in the opposite direction for 9.8%, close to REVEL, BayesDel, and AlphaMissense (72.5–76.3% and 7.3–8.2%) and better than CADD (60.5% and 13.6%) (Table S11, Figure 2A). For variants later classified as P/LP, the missense predictors contributed, on average, +1.9 to +2.0 points because they reached PP3 strong, compared with +1.3 for AVI and +0.9 for CADD. The supporting and moderate intervals met the ClinGen likelihood ratios for every predictor except CADD PP3 moderate (5.54, 95% CI 4.90–6.35, where 5.79 is required). Because the 2026 classifications could cite every predictor except AVI, the small advantage of REVEL and BayesDel may partly reflect circularity. Among the prespecified examples (Table S12), three *MYOC* (HGNC:7610) variants reclassified as likely benign by an expert panel received BP4 evidence from all five predictors. In contrast, *LDLR* (HGNC:6547) NM_000527.5:c.1381G>T p.(Gly461Cys) and *PIK3CD* (HGNC:8977) NM_005026.5:c.1002C>G p.(Asn334Lys), reclassified as likely pathogenic by expert panels, received BP4 evidence from AVI and CADD. Pathogenic variants in *PIK3CD* act by gain of function,^33^ the gene class with the highest thresholds.

**Figure 2.**
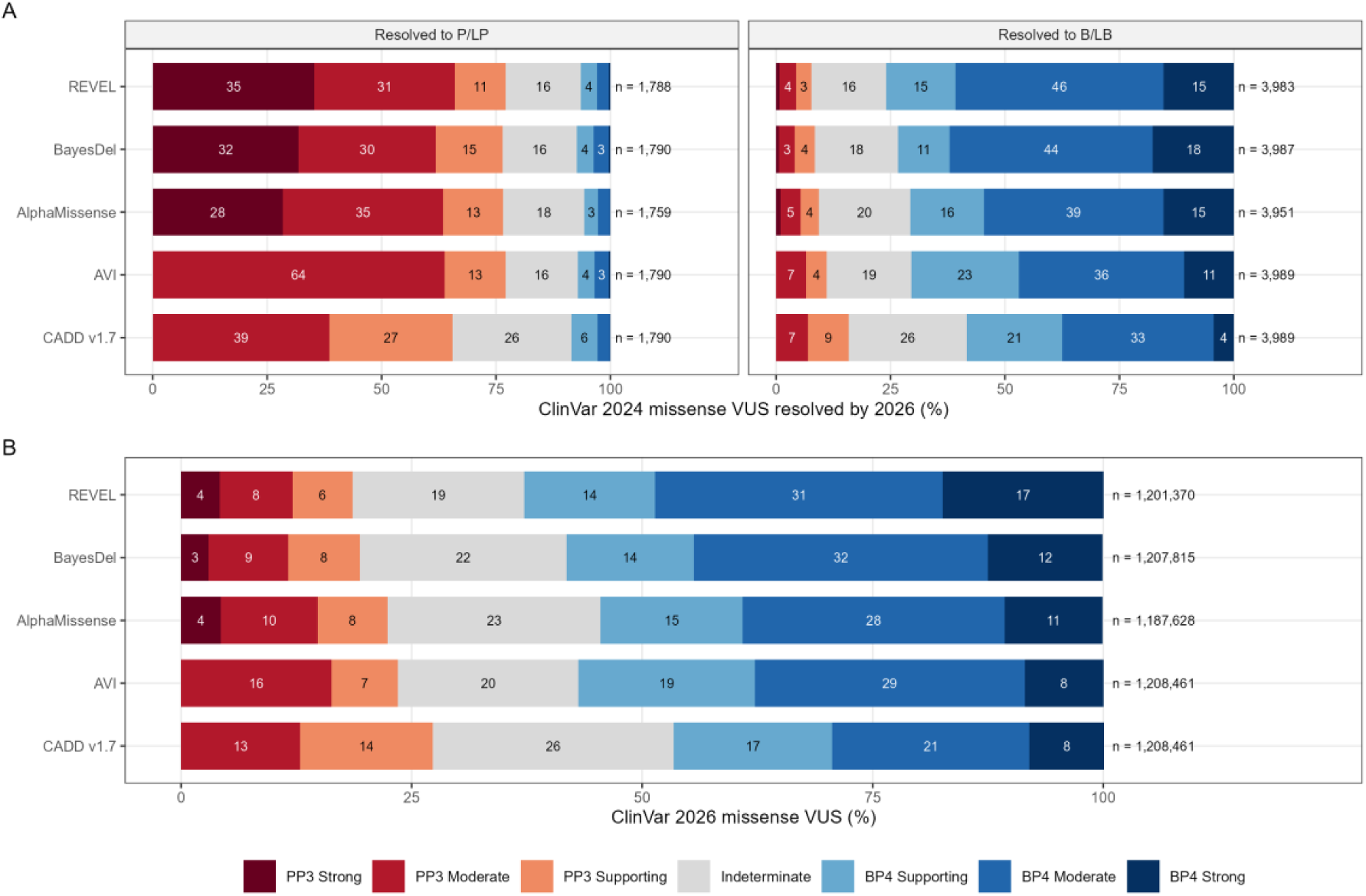
Calibrated computational evidence for missense variants of uncertain significance. (A) Missense VUS with at least one star in MANE Select transcripts of disease genes and gnomAD v4.1 allele frequency below 0.01 in the ClinVar release of January 27, 2024 that were classified P/LP (left) or B/LB (right) by the release of September 23, 2026, with the evidence that thresholds calibrated on the January 2024 release alone would have given. (B) Distribution of PP3 and BP4 evidence levels among missense VUS selected in the same way from the ClinVar release of September 23, 2026, without the allele frequency filter, for each predictor, using the thresholds calibrated on the five-predictor intersection sets (median of five seeds). Red segments are PP3 levels, blue segments are BP4 levels, and the grey segment is the indeterminate zone. Numbers inside the segments are percentages (shown when ≥ 3%), and numbers on the right give the variants scored by each predictor.

### Current VUS

Used alone, each calibrated predictor gave BP4 evidence to 46–63% and PP3 evidence to 19–27% of the 1,208,461 missense VUS of the September 2026 release, depending on the predictor (Figure 2B, Table S13). Of the 1,208,430 rare VUS, the three missense predictors and AVI agreed on pathogenic evidence for 105,497 (8.7%) and on benign evidence for 378,178 (31.3%) (Table S14). The concordant pathogenic evidence was strong for 10,553 variants, moderate for 46,931, and supporting for 48,013, and the concordant benign evidence was moderate for 167,479 and supporting for 210,699.

For 1,094 (37.1%) of the 2,948 VUS reviewed by expert panels, in 97 genes, the evidence was concordant, with PP3 strong for 186, PP3 moderate for 341, PP3 supporting for 280, BP4 supporting for 175, and BP4 moderate for 112 (Table S15, Table S16). *RUNX1* (HGNC:10471) had 217, with evidence in both directions; *RYR1* (HGNC:10483) had 101, almost all with pathogenic evidence; and the monogenic diabetes genes *HNF1A* (HGNC:11621), *HNF4A* (HGNC:5024), and *GCK* (HGNC:4195) had 137, of which 129 had pathogenic evidence. All 40 such variants in *PAH* (HGNC:8582) had pathogenic evidence, whereas benign evidence predominated in *MYOC*, *ITGA2B* (HGNC:6138), and *CDH1* (HGNC:1748).

Read with the published CADD v1.6 thresholds, the CADD v1.7 scores of 216,454 VUS (17.9%) received a different evidence level from that of the calibrated thresholds, mostly weaker benign evidence (92,776 BP4 supporting instead of moderate and 79,467 BP4 moderate instead of strong) and 30,239 PP3 supporting evidence that calibration removed. No variant changed direction between the two CADD threshold sets, but 55,003 VUS (4.6%) received evidence in opposite directions from the transferred CADD v1.6 thresholds and calibrated AVI.

### Variant classes outside missense

Most ClinVar variants outside the MANE coding sequence fell into classes dominated by one label. Only the intronic splice region (3–8 nucleotides from an exon) had enough variants of both labels to calibrate. In that class, both predictors reached PP3 strong at much lower scores than for missense variants, CADD at PHRED ≥15.45, ≥17.36, and ≥20.10 for supporting, moderate, and strong evidence, and AVI at ≥21.30, ≥22.44, and ≥24.09 (Table S17). In the other classes, the missense BP4 supporting interval was enriched for pathogenic variants in ten of eleven classes for CADD and nine for AVI, for example, among near-intronic (likelihood ratio 12.0 for CADD and 15.8 for AVI) and deep intronic variants (7.5 and 8.8), where a ratio of at most 0.42 is required (Figure 3A, Table S18). AVI discriminated better than CADD in ten of eleven classes, most clearly among near-intronic variants (AUROC 0.956 versus 0.779) (Figure 3B), and on MPRA data from 20 regulatory elements (AUROC 0.679 versus 0.565).

**Figure 3.**
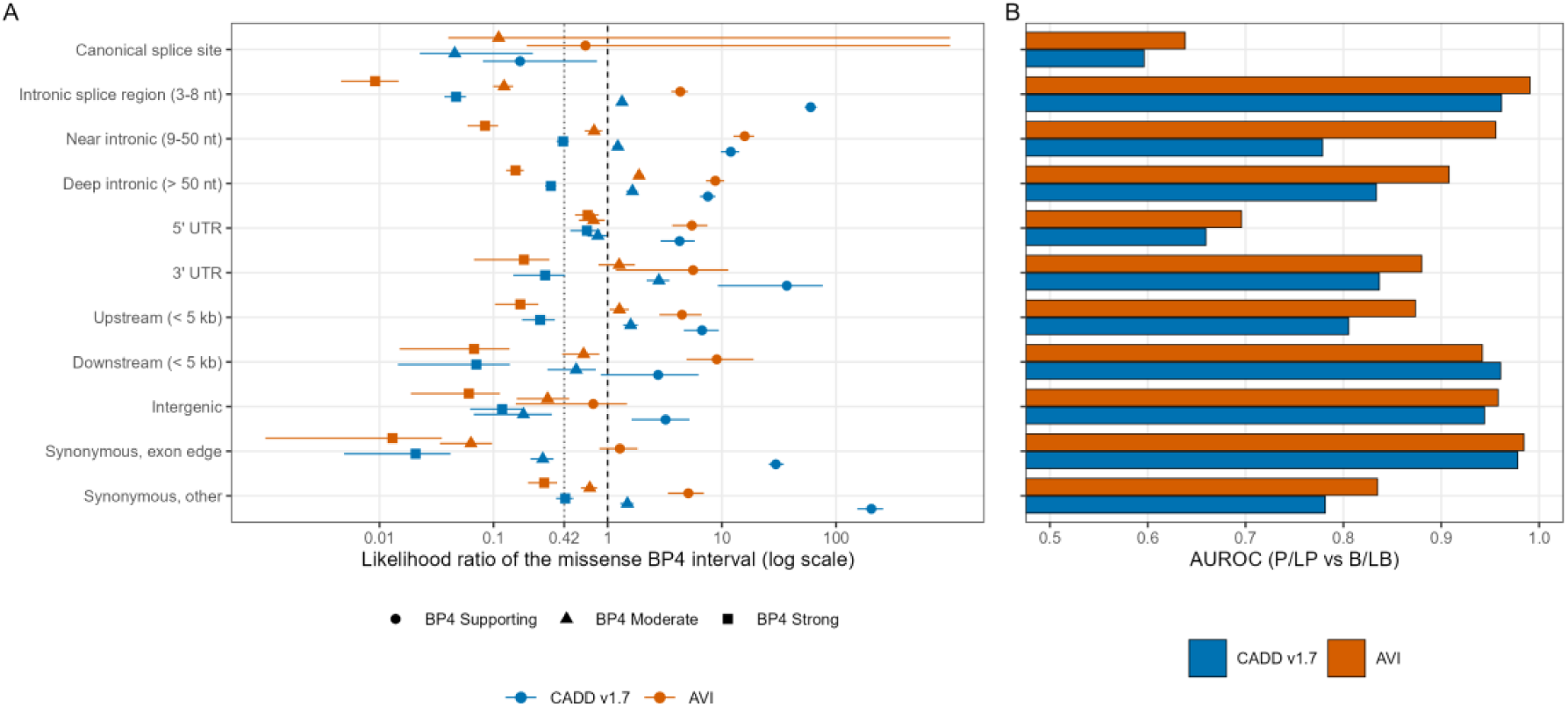
Missense-calibrated thresholds outside missense variation. (A) Likelihood ratio of each missense BP4 interval of CADD v1.7 (dark) and AVI (light), P(interval | P/LP) / P(interval | B/LB), with 95% bootstrap intervals (1,000 replicates), among ClinVar variants (release of September 23, 2026, at least one star, disease genes) outside the MANE Select coding sequence, grouped by region class. The dashed line marks a ratio of 1, and the dotted line marks the ratio of at most 0.42 that BP4 supporting evidence requires; ratios above 1 mean that the interval meant to indicate benignity is enriched for pathogenic variants. Intervals with fewer than ten variants are not shown. (B) Area under the receiver operating characteristic curve (AUROC) of each predictor for P/LP versus B/LB variants in each class. Intronic distances are measured from the nearest exon boundary.

### Transfer between model versions

On the same reference variants, CADD v1.6 and v1.7 gave similar PP3 thresholds (v1.6 ≥25.2 and ≥27.5, v1.7 ≥25.6 and ≥28.0) and discriminated equally well (AUROC 0.904 and 0.902) (Table 1, Table S19). With each version at its own thresholds, however, 38.5% of P/LP and 33.0% of B/LB variants received a different evidence level, and 868 variants moved between PP3 and BP4.

### Reference data and gene class

Our REVEL and BayesDel thresholds were less stringent than the published ones at PP3 moderate and strong and at every BP4 level.^6^ They remained so when calibrated on the December 2019 release used in the original study. Removing HGMD Public positions reduced the 2019 P/LP set from 17,330 to 5,395 variants, close to the 5,704 of the original study, and restored the published thresholds (Table S20), whereas the CADD v1.7 and AVI supporting and moderate thresholds changed by 0.5 units or less. Across ClinVar releases, PP3 moderate and strong thresholds relaxed modestly for REVEL, BayesDel, and CADD, the predictors recommended in 2022, but not for AlphaMissense or AVI (Table S21), a pattern consistent with circularity.

Gene class also changed the thresholds, with the same pattern for all five predictors (Figure S6, Table S22). Constrained, dominant, and gain-of-function or dominant-negative genes required higher PP3 scores and allowed BP4 at higher scores than tolerant or recessive genes. Class-specific thresholds would change the evidence level for 3–8% of variants in loss-of-function genes, 26–44% in tolerant genes, and 17–54% in gain-of-function or dominant-negative genes.

## DISCUSSION

Genome-wide predictors provide reliable computational evidence only when their thresholds match the variant class, the model version, and the reference data from which they were derived. CADD v1.7 and AVI reached moderate pathogenic and strong benign evidence for rare missense variants, and their thresholds held both for variants classified after calibration and for most VUS resolved since January 2024. Table 2 turns these results into recommendations for clinical laboratories.

**Table 2.**
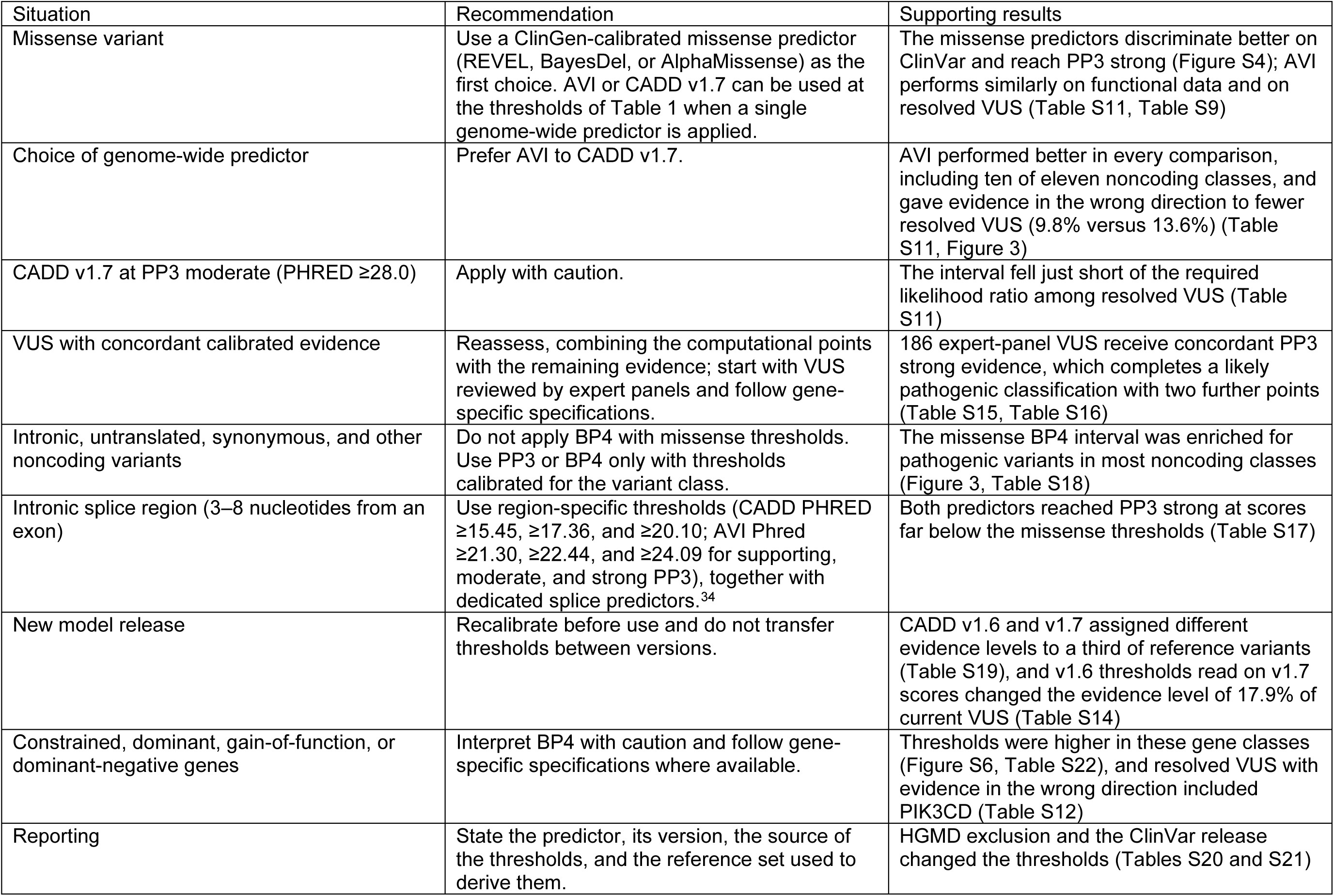
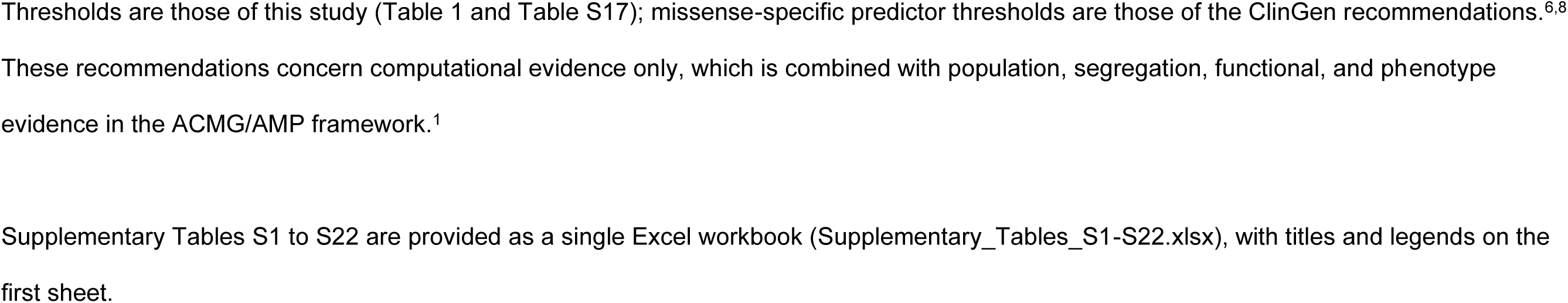
Recommendations for using genome-wide predictors as PP3/BP4 evidence.

Among the genome-wide predictors, AVI performed most consistently, scoring higher than CADD v1.7 on ClinVar, on variants classified after calibration, on resolved VUS, on functional data, and in ten of eleven noncoding classes, although the size of the difference varied. For missense variants, it matched REVEL and BayesDel on functional data and narrowly missed PP3 strong. Because no ClinVar classification could have used it, its results are free of the circularity that may favor older predictors. For a laboratory that needs one genome-wide predictor, these results favor AVI over CADD.

For missense variants, REVEL, BayesDel, and AlphaMissense remain the first choice, because they perform better on ClinVar and reach PP3 strong, in line with earlier benchmarks.^6,8,17^ Fewer than 1% of missense VUS were resolved in two and a half years; most were benign, consistent with a large clinical cohort.^3^ For these variants, calibrated evidence pointed in the final direction for about three-quarters of cases and added one to two ACMG/AMP points on average, which moves a variant across a classification boundary only when the other evidence already places it near one. It pointed in the wrong direction for 7-14%, including variants in gain-of-function genes. Calibrated predictors still err, but each point they contribute comes with a known likelihood ratio.

To judge the practical consequences, we examined the VUS that remain unresolved today. Concordant PP3 strong evidence, which 10,553 rare missense VUS receive, completes a likely pathogenic classification with two further points, for example, two supporting criteria, and concordant BP4 moderate evidence, which 167,479 receive, completes a likely benign classification unless the remaining evidence adds two or more pathogenic points. These numbers describe what the computational evidence allows, not what the classifications will become, because population, segregation, functional, and phenotypic evidence are not recorded as points in ClinVar. The list of expert-panel VUS (Tables S15 and S16) is a practical starting point for reassessment, although the panels may already have applied their own computational criteria.^7^ It concentrates on *RUNX1*, *RYR1*, the monogenic diabetes genes, *LDLR*, and *PAH*. *RUNX1*, *RYR1*, *HNF1A*, and *HNF4A* are among the most constrained genes (LOEUF deciles 0–2), which required higher PP3 scores in our gene-class analysis, so PP3 evidence in these genes should be weighed against the gene-specific specifications.

Variants outside the missense class revealed the main limitation of applying missense-derived thresholds to other variant classes. One CADD cutoff applied to every variant type gives false benign evidence to pathogenic intronic, untranslated, and synonymous variants, which score lower than pathogenic missense variants, and withholds pathogenic evidence from intronic splice-region variants, for which both predictors reached PP3 strong at scores well below the missense thresholds. ClinGen recommends dedicated splice predictors and specific rules for splicing.^34^ Genome-wide scores should provide PP3 or BP4 evidence for noncoding variants only with thresholds calibrated for each variant class, and BP4 should not be applied to noncoding variants with missense thresholds.

Thresholds are also tied to the model version and the reference data. CADD v1.7 did not discriminate missense variants better than v1.6, yet a third of reference variants received a different evidence level, because retraining reorders variants by genome-wide rank. Reusing the v1.6 thresholds with v1.7 scores, the shortcut a software update invites, would change the evidence level of 18% of current VUS, mostly by weakening benign evidence. Laboratories should therefore recalibrate when they change model release, and interpretation software should state which version and thresholds it applies. Our REVEL and BayesDel thresholds matched the published values only after HGMD-listed positions were removed, so a laboratory that recalibrates without HGMD Professional should expect less stringent BP4 thresholds, and thresholds built on different reference sets should not be mixed. PP3 thresholds drifted across ClinVar releases for the predictors recommended in 2022 but not for AVI, suggesting that predictor evidence in ClinVar submissions is beginning to shape the labels used to recalibrate those predictors.^12^ Temporal hold-out sets and negative controls such as AVI can protect future recalibrations.

All five predictors also required higher PP3 scores in constrained, dominant, and gain-of-function or dominant-negative genes, which may reflect a different prior probability of pathogenicity or gene-level constraint partly encoded in the scores. Class-specific thresholds or priors, which would change the evidence level of up to half of the variants in some classes, deserve formal study alongside the gene-specific specifications of expert panels.^7^

Calibrated computational evidence is weighed alongside frequency criteria, and a variant that is common in a population rarely causes a rare Mendelian disease.^1,35,36^ Reference databases still represent mainly European populations,^37^ and individuals of non-European genetic ancestry are more likely to carry VUS.^3,37^ Latin American resources such as the Mexico City Prospective Study, the Mexican Biobank, admixed Brazilian genomes,^38–40^ and the 1000 Genomas Para Chile project can supply the ancestry-specific frequencies that BA1, BS1, and PM2 require (Figure S7); how they interact with calibrated thresholds will be examined separately.

Our study has limitations. The labels come from ClinVar, which may already incorporate computational evidence, and the negative-control predictor, the temporal hold-out, the resolved VUS, and the functional data reduce but do not remove this circularity. We could not exclude variants from HGMD Professional or predictor training sets, and could not replicate the gnomAD quality filters used in the original study. The functional benchmark covered 39 genes with assays of varying relevance to disease; the gene-class analyses used a single prior; and, aside from missense variation, only the intronic splice region could be calibrated, using the missense prior and one seed.

In summary, CADD v1.7 and AVI can provide calibrated PP3 and BP4 evidence for missense variants, and AVI performed better in most of the settings we examined, inside and outside the coding sequence.

The thresholds depend on context and should be revisited whenever the variant class, the reference data, or the model version changes, and laboratories should report the source of every threshold they apply.

The expert-panel VUS with concordant evidence listed here are a reasonable starting point for reassessment.

### Data and code availability

The analysis plan and its amendments, every executed script with its SHA-256 hash, console logs, figure source data, and a renv lockfile are available at https://github.com/JPMirandaM/cadd-alphagenome-acmg-calibration (archived at https://doi.org/10.5281/zenodo.23790410), together with the 483,690 ClinVar missense VUS with concordant calibrated evidence (calibration/results/v17_avi/vus_candidates/vus_concordant_evidence.tsv.gz). The ClinGen Sequence Variant Interpretation calibration code that we ported is available at https://github.com/vpejaver/clingen-svi-comp_calibration. Third-party data are not redistributed, and their sources and terms are listed in THIRD_PARTY_NOTICES.md and executed_pipeline/data_manifest.tsv. A genome-wide catalog of SNVs with PP3-level scores will be described separately, subject to the score providers’ license terms.

## Acknowledgments

We thank the ClinGen Sequence Variant Interpretation working group for making their calibration code and data publicly available; the developers of CADD, AlphaGenome, REVEL, BayesDel, and AlphaMissense for releasing their scores; the submitters to ClinVar; the gnomAD consortium; MaveDB, and the authors of the MAVE score sets used in this study.

## Funding

This work was funded by Bupa Chile through the 1000 Genomas Para Chile project. The funder had no role in the design of the study, the collection and analysis of data, or the decision to publish.

## Author contributions

Conceptualization: J.P.M.; Data curation: J.P.M.; Formal analysis: J.P.M.; Investigation: J.P.M.; Methodology: J.P.M.; Software: J.P.M.; Visualization: J.P.M.; Writing-original draft: J.P.M.; Writing-review & editing: J.P.M., L.C., I.A., P.A., C.S.; Funding acquisition: J.P.M., C.S.; Project administration: J.P.M., C.S.; Resources: J.P.M.; Supervision: J.P.M.; Validation: J.P.M., P.A.

## Ethics Declaration

This study used only public data: predictor scores, ClinVar variant classifications, summary-level gnomAD allele frequencies, and published functional assay data. No individual-level human data were accessed, no human participants were enrolled, and no biological samples were obtained or analyzed. All data were already de-identified and aggregated at source. The study therefore did not qualify for submission to an Institutional Review Board or Research Ethics Committee in the authors’ jurisdiction, and informed consent was not required.

## Conflict of Interest

The authors declare no conflict of interest. All authors are employed by Bupa Chile, which funded this work through the 1000 Genomas Para Chile project.

## Declaration of AI and AI-assisted technologies in the writing process

During the preparation of this work, the authors used Claude Code (Anthropic) in order to write and run analysis code under the authors’ direction and to draft and edit the manuscript text. After using this tool, the authors reviewed and edited the content as needed and took full responsibility for the content of the publication. The authors designed the study, defined the analysis plan and any amendments before each analysis, and reviewed the code, results, and text.

## Notes

https://github.com/JPMirandaM/cadd-alphagenome-acmg-calibration

https://doi.org/10.5281/zenodo.23790410

### Competing Interest Statement

The authors have declared no competing interest.

### Summary of Updates

This version reports no new analysis and changes no result. Every count, threshold, likelihood ratio, AUROC and conclusion is identical to version 1. The manuscript was reformatted for a different target journal and corrected on points of nomenclature and citation style. Abstract: restructured into Purpose, Methods, Results and Conclusion, and shortened from 250 to 198 words. Length: the body was shortened from 4054 to 3970 words, mainly by condensing the Methods, whose detail moved to the Supplemental Methods. Display items: reduced from nine to five. Table 2 and Table 3 of version 1 are now Supplementary Tables S11 and S15, and Figure 2 and Figure 5 are now Supplementary Figures S4 and S6. Table 4 of version 1 is now Table 2, Figure 3 is now Figure 2, and Figure 4 is now Figure 3. The supplementary tables were renumbered from S1 to S20 into S1 to S22. Figures: redrawn in a colour-blind-safe palette with no red and green pairing. Version 1 was grayscale. References: reduced to 40 entries, cited as superscript numerals and numbered by order of first appearance. Citation style was made consistent across the tables, where some citations were still bracketed and one pointed to a number that no longer existed after the list was shortened. Nomenclature: HGNC gene identifiers were added at the first mention of each gene. The two variants named in the text are now given in full on the MANE Select transcript and on GRCh38. The PIK3CD variant p.Asn334Lys is now identified by its coding change on transcript NM_005026.5, because ClinVar holds two distinct single nucleotide variants that produce that same substitution and only one of them was scored here. Every variant description in the manuscript and in the supplementary tables was verified with VariantValidator. Other changes: the statement that the AlphaGenome Variant Impact score had not yet been calibrated was removed, because a calibration is now reported here. The source of that score is cited as a preprint with its DOI. A pointer to the ported ClinGen Sequence Variant Interpretation calibration code was added. Wording about ancestry was changed from patients to individuals of non-European genetic ancestry. Data and code availability is unchanged, including the archived Zenodo record.

