## Supplemental information for "Beyond a single threshold: calibration of CADD v1.7 and AlphaGenome for ACMG/AMP computational evidence across variant classes, model versions, and reference data"

This document contains the Supplemental Methods, Supplemental Figures S1 to S7, and the titles and legends of Supplemental Tables S1 to S22, which are provided in the Excel workbook Supplementary_Tables_S1-S22.xlsx, one sheet per table. Scripts, logs and source data are available in the study repository https://github.com/JPMirandaM/cadd-alphagenome-acmg-calibration (archived at https://doi.org/10.5281/zenodo.23790410).

### Supplemental methods

### Analysis plan and amendments

The analysis of threshold drift across ClinVar releases was prompted by differences between our REVEL and BayesDel thresholds and the published ones, and the HGMD exclusion followed from the drift analysis. The analyses of VUS resolved after calibration and of current VUS were registered before any of their results were seen. All amendments are listed with their dates in Table S1.

### Predictors and annotation

CADD v1.7 GRCh38 PHRED scores for all 8,812,917,339 possible SNVs were checked against the published MD5 checksum. BayesDel scores without allele frequency were lifted from hg19 to GRCh38 and kept only where the hg19 reference base matched. When a predictor yielded multiple scores for a single SNV, the maximum was used. We enumerated all 100,316,172 possible SNVs in the coding sequences of the 19,229 GENCODE v50 MANE Select transcripts, and their annotated consequences agreed with the ClinVar molecular consequence for 99.99% of 3,405,151 ClinVar SNVs.

### Reference sets

Labeled variants were missense in the MANE Select transcript and not stop-gained, stop-lost, or start-lost in any overlapping MANE transcript, and their gnomAD v4.1 exome and genome allele frequencies were both below 0.01. Three deviations from the original study were prespecified. HGMD-listed variants were kept in the main analysis because their effect was tested separately, training variants were kept because neither CADD nor AVI is trained on ClinVar labels, and the gnomAD quality filters of the original study could not be applied because they are not available in the Ensembl VEP cache.

### Calibration and sensitivity analyses

The R implementation keeps the window arithmetic, inclusive window bounds, and threshold rules of the original MATLAB code. The threshold for each strength is counted from the end of the scale. Labeled and unlabeled variants were resampled independently in each bootstrap replicate. On the original authors' deposited data, the code reproduced the published CADD v1.6 thresholds (PP3 25.3 and 28.1; BP4 22.7 and 17.3). The four sensitivity analyses (Table S5) required at least two review stars, defined rarity as allele frequency below 0.001, resampled genes rather than variants, and excluded missense SNVs within three nucleotides of an internal exon boundary.

### Resolved and current VUS

Resolved VUS were analyzed for all five predictors across variants scored by all five predictors, and the primary analysis was restricted to variants with gnomAD allele frequency below 0.01. The likelihood ratio for each interval among resolved variants (P/LP versus B/LB) was estimated using 2,000 bootstrap replicates. Illustrative examples (Table S11) were selected using a fixed rule: resolved variants whose 2026 record was reviewed by an expert panel or practice guideline, up to three per direction, with the highest AVI points. For current VUS, variants reviewed by an expert panel or practice guideline are listed individually (Table S14), and the counts of variants whose evidence level or direction differed between transferred CADD v1.6 thresholds and calibrated CADD v1.7 and AVI are given in Table S13.

### ClinVar release series

The September 2026 reference set was split into variants already present in November 2022, and variants added later. AVI was a prespecified negative control, so that a relaxation of PP3 thresholds in later releases for REVEL, BayesDel, and CADD, but not for AVI, would suggest circularity.

### Functional assays, gene classes, and variant classes

MaveDB score sets were mapped to GRCh38 with the MaveDB mappings. Protein-level sets were kept when at least 90% of mapped reference residues matched the MANE protein, and variants classified discordantly across sets were discarded. Likelihood ratios were estimated with 2,000 bootstrap replicates, and the AUROC was computed overall and per gene. Mode of inheritance was taken from ClinGen gene–disease validity curations, and from the Gene Curation Coalition when ClinGen had no curation. CADD v1.6 scores for the 2,093,102 reference SNVs were obtained from the official v1.6 file.

### Software

Bioconductor packages included Rsamtools, GenomicRanges, rtracklayer, and Biostrings.

### Supplemental figures

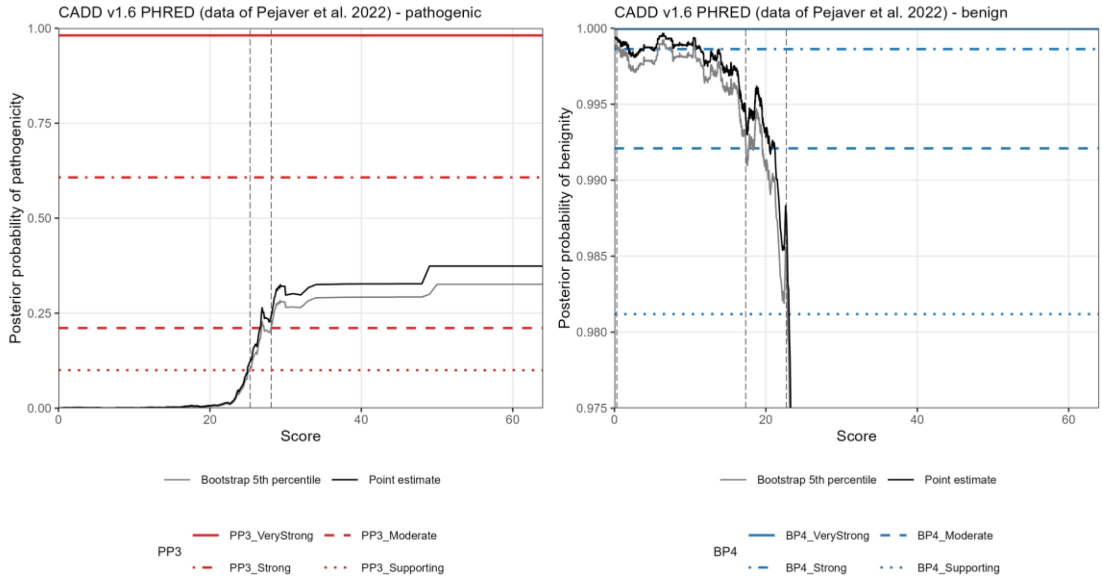


**Figure S1. Reproduction of the published CADD v1.6 calibration.** Local posterior probability of pathogenicity and benignity for CADD v1.6 PHRED computed with our implementation on the data of Pejaver et al. (2022), with 10,000 bootstrap replicates (seed 20260925). Lines as in Figure 1.

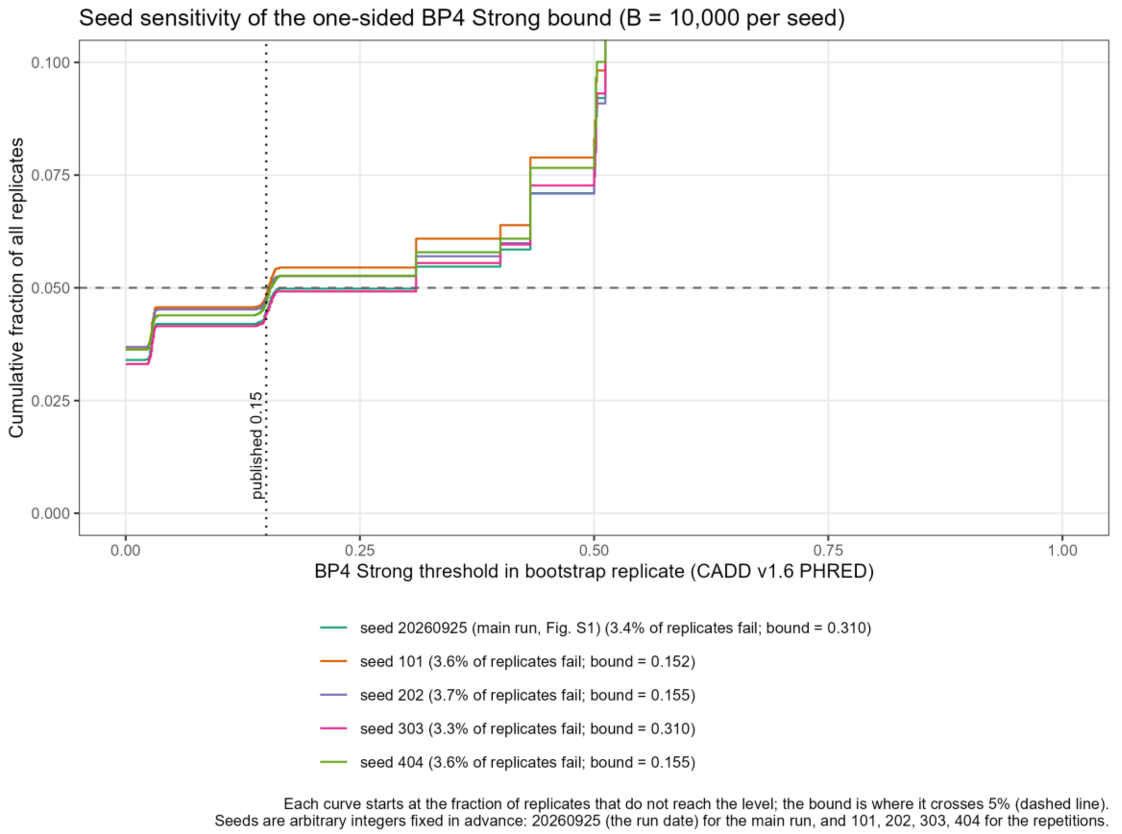


**Figure S2. Seed dependence of the CADD v1.6 BP4 strong threshold.** Cumulative distribution of the 10,000 bootstrap thresholds for five random seeds, with replicates that fail to reach the level counted first. The one-sided 95% bound falls between two plateaus of the distribution, which explains the alternation between 0.15 and 0.31.

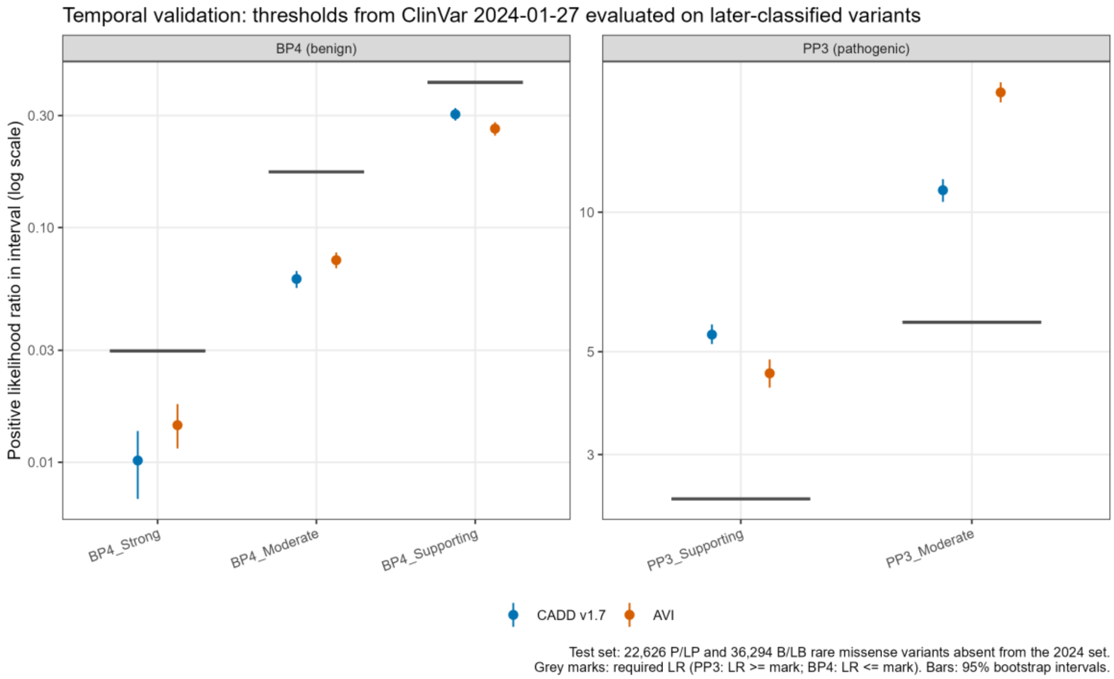


**Figure S3. Temporal validation.** Positive likelihood ratios, with 95% bootstrap intervals, of each evidence interval of CADD v1.7 and AVI calibrated on the ClinVar release of January 27, 2024 and evaluated on the 58,920 variants classified afterwards. Horizontal lines mark the likelihood ratios required for each evidence strength.

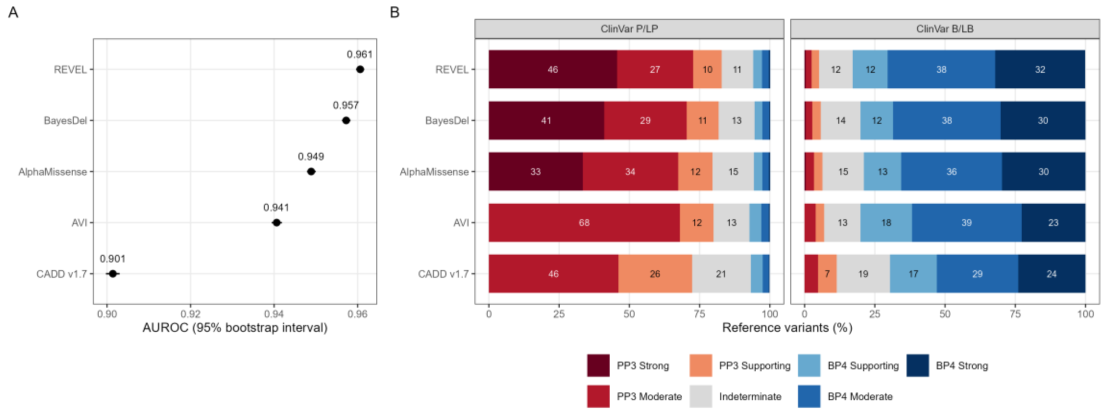


**Figure S4. Discrimination and evidence levels of five predictors on ClinVar.**

(A) Area under the receiver operating characteristic curve (AUROC) with 95% bootstrap intervals (1,000 replicates) for separating P/LP from B/LB variants. (B) Percentage of P/LP (left) and B/LB (right) reference variants assigned to each evidence level by each predictor's calibrated thresholds (median of five seeds), computed on the calibration set itself. Each bar adds up to 100%. Red segments on the left of each bar are PP3 levels, blue segments on the right are BP4 levels, and the grey segment in between is the indeterminate zone, in which scores fall between the PP3 supporting and BP4 supporting thresholds and provide no computational evidence. A well-calibrated and informative predictor places most P/LP variants in red segments and most B/LB variants in blue segments, with a narrow grey zone. Numbers within the segments are percentages (shown when ≥6%). The reference sets are the 2026 ClinVar variants (P/LP versus B/LB, at least one star, rare MANE missense) and the gnomAD variants scored by all five predictors (134,340 labeled variants).

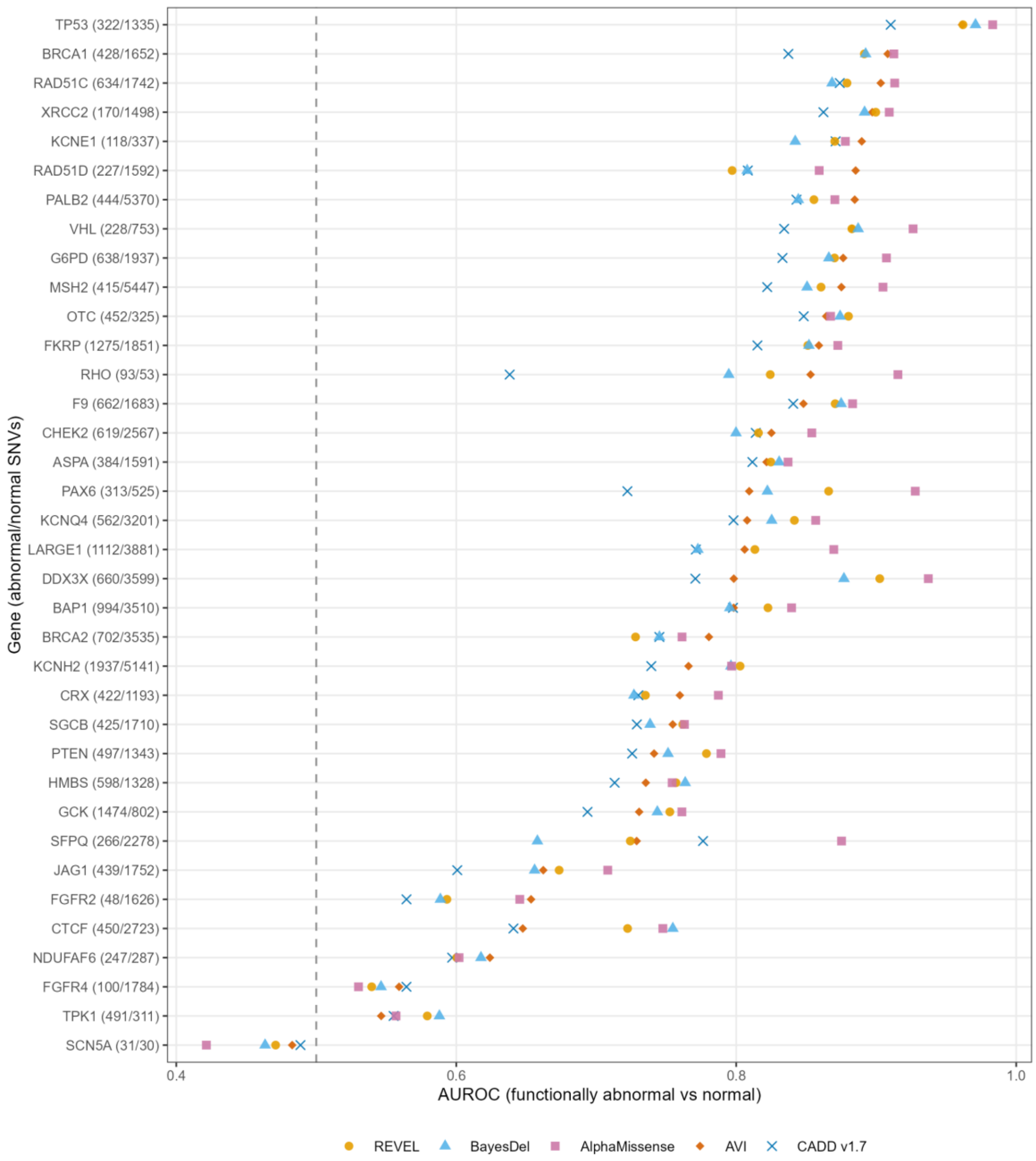


**Figure S5. Discrimination of MAVE functional classes by gene.** AUROC of five predictors for separating functionally abnormal from normal MANE missense SNVs in each of the 36 genes with at least 30 abnormal and 30 normal variants, from MaveDB score sets with normal and abnormal functional classes. Numbers in parentheses give abnormal and normal SNVs. Genes are ordered by the AUROC of AVI, and the dashed line marks an AUROC of 0.5.

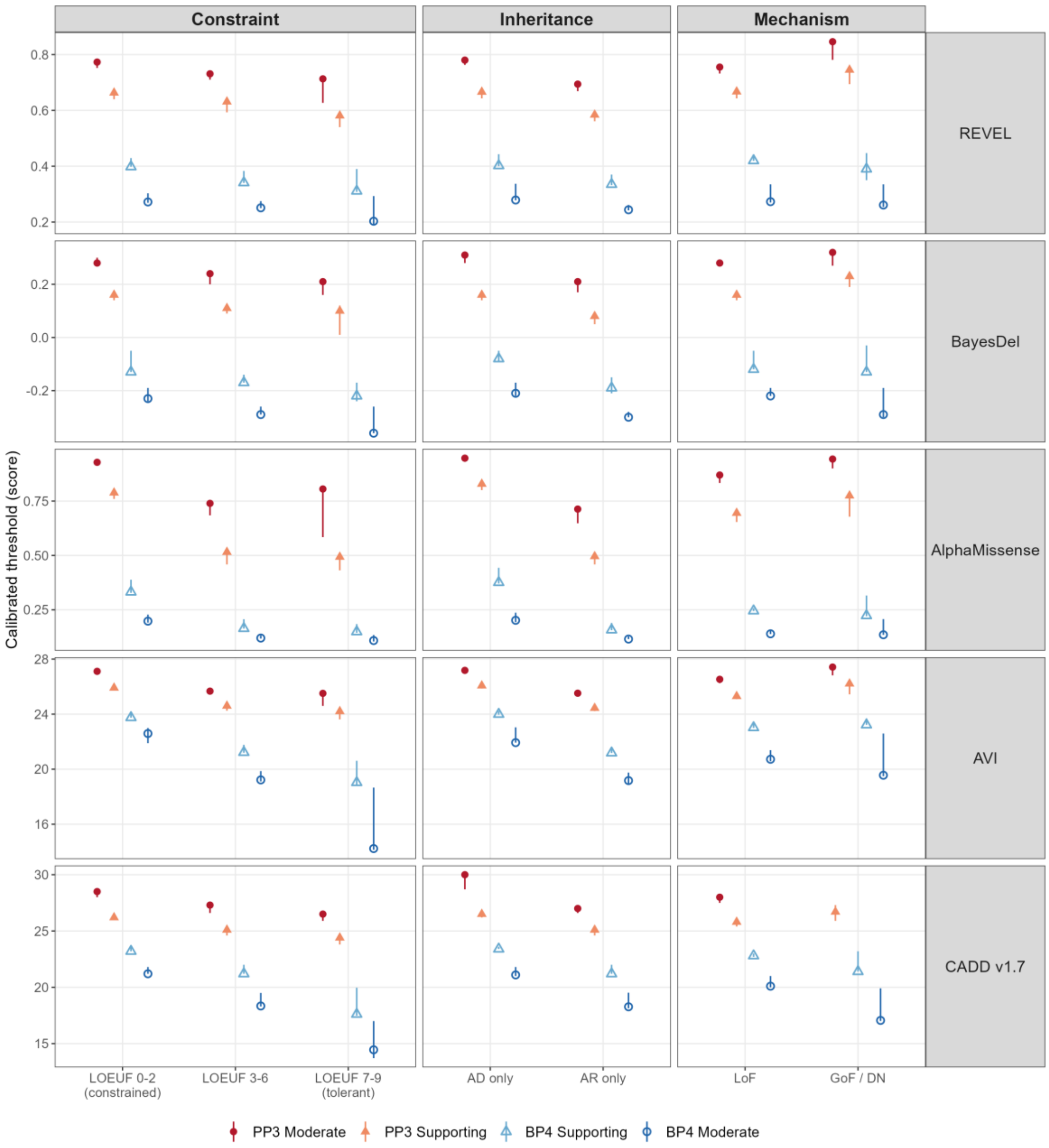


**Figure S6. Calibrated PP3 and BP4 thresholds by gene class.**

Thresholds of five predictors (rows) in gene classes defined by constraint (gnomAD v4.1 LOEUF deciles 0–2, 3–6, and 7–9), mode of inheritance (autosomal dominant only or autosomal recessive only), and disease mechanism (loss of function, LoF, or gain of function and dominant negative, GoF/DN). Filled symbols are PP3 thresholds and open symbols are BP4 thresholds, with circles for moderate and triangles for supporting evidence. Symbols mark the final thresholds (one-sided 95% bootstrap bound) and bars span the 2.5th to 97.5th percentiles of 10,000 bootstrap thresholds. Each class was calibrated on the 2026 ClinVar and gnomAD intersection sets, restricted to that class's genes, using a global prior of 0.0441. Missing points indicate levels that were not reached.

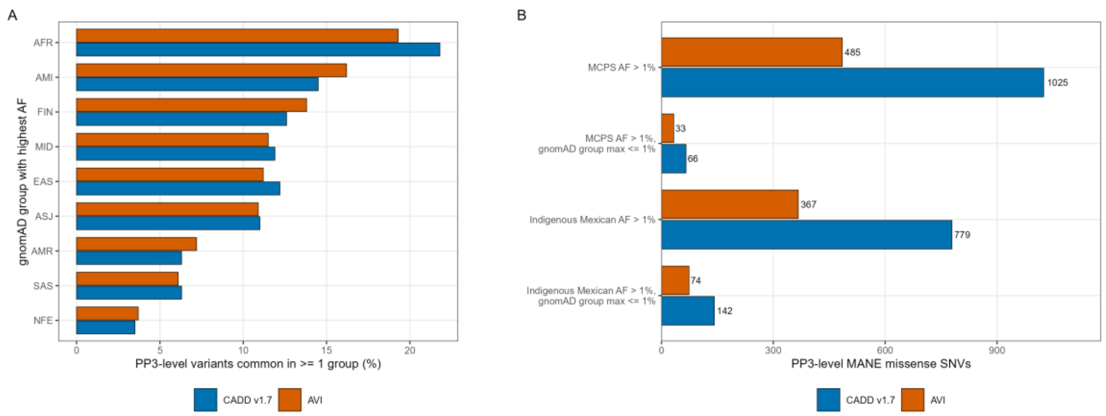


**Figure S7. Population frequency of missense variants with PP3-level scores.**

(A) Among MANE missense SNVs that reach a PP3 level (supporting or moderate, main-analysis median thresholds) and are common (AF > 1%) in at least one gnomAD v4.1 genetic ancestry group, the percentage for which each group has the highest frequency. Group AF is the maximum of the exome and genome AF of the group. AFR, African/African American; AMI, Amish; AMR, admixed American; ASJ, Ashkenazi Jewish; EAS, East Asian; FIN, Finnish; MID, Middle Eastern; NFE, non-Finnish European; SAS, South Asian. (B) Number of PP3-level MANE missense SNVs with AF > 1% in the Mexico City Prospective Study (MCPS), overall and in the Indigenous Mexican ancestry component, and the number of these that are at or below 1% across all gnomAD groups. MCPS frequencies are from the RGC-MCPS release 20230130 (Ziyatdinov et al. 2023); data provided by the Regeneron Genetics Center and collaborators.

### Supplemental tables

**Table S1. Prespecified analysis plan and dated amendments.**

The analysis plan was finalized on 25 September 2026 before any CADD v1.7 or AVI calibration results were obtained and tagged in the version-controlled repository (analysis-plan-v1). Each later analysis was specified in a dated amendment committed before it was run. The Timing column indicates whether the amendment was motivated by results already observed, and the last column indicates whether the analysis is reported in this article. Post hoc analyses that were not specified in any amendment are listed at the end.

*Provided in Supplementary_Tables_S1-S22.xlsx, sheet S1 (17 rows).*

**Table S2. Predictors, score precision and window increments.**

Score sources and versions, the precision at which scores were used and the window increment of the local posterior method, chosen to match the original study.

*Provided in Supplementary_Tables_S1-S22.xlsx, sheet S2 (6 rows).*

**Table S3. Published versus reproduced CADD v1.6 thresholds.**

Thresholds of Pejaver et al. (2022, Table 2) and those obtained with our R implementation on the authors' deposited data (10,000 bootstrap replicates, seed 20260925).

*Provided in Supplementary_Tables_S1-S22.xlsx, sheet S3 (8 rows).*

**Table S4. Seed sensitivity of the CADD v1.6 reproduction.**

Thresholds for five random seeds (10,000 replicates each) with the fraction of bootstrap replicates that did not reach each level.

*Provided in Supplementary_Tables_S1-S22.xlsx, sheet S4 (20 rows).*

**Table S5. Sensitivity analyses of the CADD v1.7 and AVI calibration.**

Thresholds (seed 20260925) in the main analysis and in the four prespecified sensitivity analyses, namely review status of at least two stars (two_stars), rarity defined as AF below 0.001 (AF_below_0.001), gene-level bootstrap (gene_bootstrap), and exclusion of missense SNVs within 3 nucleotides of an internal exon boundary (no_exon_edge_3nt). Empty cells indicate levels not reached.

*Provided in Supplementary_Tables_S1-S22.xlsx, sheet S5 (16 rows).*

**Table S6. Temporal hold-out validation.**

Positive likelihood ratios of each evidence interval of CADD v1.7 and AVI calibrated on the ClinVar release of 27 January 2024 and evaluated on 58,920 variants (22,626 P/LP, 36,294 B/LB) first classified afterward, with 95% bootstrap intervals (1,000 replicates), the required ratio, and whether it was met at the point estimate and at the conservative 95% bound. The second block compares 2024 and 2026 thresholds.

*Provided in Supplementary_Tables_S1-S22.xlsx, sheet S6 (20 rows).*

**Table S7. Calibration and discrimination of five predictors on the intersection set.**

Thresholds (median [range] over five seeds) and AUROC and AUPRC with 95% bootstrap intervals and paired differences versus AVI (1,000 replicates), on the 59034 P/LP and 75306 B/LB ClinVar variants (4725 genes) and 1888558 gnomAD variants scored by all five predictors.

*Provided in Supplementary_Tables_S1-S22.xlsx, sheet S7 (25 rows).*

**Table S8. MaveDB score sets used for validation.**

The 89 MaveDB score sets include normal and abnormal functional classes; gene, calibration used, mapping level, number of scored and classified variants, protein reference-residue match rate, license, and primary publication (PubMed ID). The second block provides classified MANE missense SNVs per gene after excluding discordant classifications. Please cite MaveDB and the primary publication of each score set.

*Provided in Supplementary_Tables_S1-S22.xlsx, sheet S8 (128 rows).*

**Table S9. Calibrated evidence intervals on MAVE functional classes.**

For each predictor and ClinVar-calibrated evidence interval, the likelihood ratio P(interval | functionally abnormal) / P(interval | functionally normal) with 95% bootstrap interval (2,000 replicates) and abnormal/normal variants in the interval, and the pooled AUROC, on 93,154 MANE missense SNVs in 39 genes (18,936 abnormal, 74,218 normal) from 89 MaveDB score sets. Functional abnormality is not pathogenicity; the ratios test direction and ordering, not the ClinGen targets.

*Provided in Supplementary_Tables_S1-S22.xlsx, sheet S9 (8 rows).*

**Table S10. Calibrated evidence for ClinVar VUS of January 2024 that were resolved by September 2026.**

Amendment A15. VUS (>= 1 star, MANE Select missense, disease gene) in the ClinVar release of 27 January 2024, followed to the release of 23 September 2026. Thresholds: r2024 = calibrated on the January 2024 release only (five-predictor intersection, primary); main2026 and inter2026 = 2026 thresholds (secondary). Blocks: set sizes; percentage of variants with PP3 or BP4 evidence and mean ACMG/AMP points by 2026 outcome; concordance of the evidence with the 2026 classification; likelihood ratio of each interval, P(interval | resolved P/LP) / P(interval | resolved B/LB), with 95% bootstrap interval (2,000 replicates), the ratio required and whether the conservative bound meets it. Subset AF<0.01 (gnomAD v4.1 group maximum) is primary.

*Provided in Supplementary_Tables_S1-S22.xlsx, sheet S10 (297 rows).*

**Table S11. Calibrated computational evidence for ClinVar missense VUS of January 2024 that were resolved by September 2026.**

Each interval cell gives the likelihood ratio P(interval | later classified P/LP) / P(interval | later classified B/LB) [95% bootstrap interval, 2,000 replicates] and, in parentheses, variants later classified P/LP / B/LB in the interval. Variants were VUS with at least one star, MANE Select missense in disease genes, and gnomAD v4.1 allele frequency below 0.01 in the ClinVar release of January 27, 2024 (1,790 later classified P/LP and 3,989 B/LB in the release of September 23, 2026). Evidence intervals use thresholds calibrated only to the January 2024 release for the five predictors across the variants scored by all of them. Mean points follow the ACMG/AMP point system (±1 supporting, ±2 moderate, ±4 strong; Tavtigian et al. 2020). —, level not reached. *The conservative bound of the 95% interval does not meet the likelihood ratio that the strength requires (5.79 for PP3 moderate, 33.53 for PP3 strong, 1/33.53 for BP4 strong). Results with 2026 thresholds and for all variants are given in Table S10.

*Provided in Supplementary_Tables_S1-S22.xlsx, sheet S11 (10 rows).*

**Table S12. Illustrative resolved VUS.**

Variants selected by the rule fixed in amendment A15 before inspection: VUS in January 2024 whose September 2026 record was reviewed by an expert panel or practice guideline, up to three per direction with the largest absolute AVI points (r2024 thresholds), ties broken by genomic position. Scores and evidence levels of all five predictors with the r2024 thresholds.

*Provided in Supplementary_Tables_S1-S22.xlsx, sheet S12 (6 rows).*

**Table S13. Computational evidence for ClinVar missense variants of uncertain significance.**

Number and percentage of 1,208,461 ClinVar missense VUS (at least one star, MANE Select, disease genes) at each evidence level for each predictor, with ACMG/AMP points, using main-analysis thresholds (CADD, AVI) or intersection thresholds (all five), for all VUS and for VUS with AF below 0.01; the second block cross-tabulates CADD and AVI levels.

*Provided in Supplementary_Tables_S1-S22.xlsx, sheet S13 (161 rows).*

**Table S14. Concordant calibrated evidence and transferred thresholds for all ClinVar missense VUS.**

Amendment A16. The 1,208,430 rare (gnomAD v4.1 AF max < 0.01) MANE Select missense VUS with at least one star in disease genes, ClinVar release of 23 September 2026. Blocks: thresholds used (CADD and AVI, Table 1; REVEL, BayesDel and AlphaMissense, published ClinGen values; CADD16, published CADD v1.6 values applied to CADD v1.7 scores); number of VUS by concordant ACMG/AMP points, for all VUS and for those reviewed by an expert panel or practice guideline (3-4 stars), and number whose evidence level or direction differs between transferred CADD v1.6 thresholds and calibrated CADD v1.7 or AVI thresholds; cross-tabulations of evidence levels under transferred CADD v1.6 thresholds against calibrated CADD v1.7 and against calibrated AVI.

*Provided in Supplementary_Tables_S1-S22.xlsx, sheet S14 (83 rows).*

**Table S15. ClinVar missense VUS reviewed by expert panels with concordant calibrated computational evidence, by gene.**

The 23 genes with at least ten such variants (856 of 1,094) are shown. Expert-panel VUS, rare (gnomAD v4.1 allele frequency below 0.01) MANE Select missense VUS in the ClinVar release of September 23, 2026, reviewed by an expert panel or practice guideline. Concordant, VUS for which REVEL, BayesDel, and AlphaMissense (published ClinGen thresholds of Pejaver et al. 2022 and Bergquist et al. 2025) and AVI (Table 1) all gave evidence in the same direction; the level is the weakest among the three missense predictors. Under the ACMG/AMP point system of Tavtigian et al. (2020), PP3 strong (+4 points) completes a likely pathogenic classification (≥6 points) when the remaining evidence adds two points, PP3 moderate when it adds four, and BP4 moderate (−2 points) a likely benign classification (≤−1 point) unless the remaining evidence adds two or more points. Expert panels may already have applied genespecific computational criteria, so these are candidates for reassessment rather than reclassification. Examples are the first three PP3 strong variants by genomic position (protein change in the MANE Select transcript). All 1,094 variants, with scores and evidence from each predictor, are listed in Table S16.

*Provided in Supplementary_Tables_S1-S22.xlsx, sheet S15 (23 rows).*

**Table S16. ClinVar missense VUS reviewed by an expert panel with concordant calibrated computational evidence.**

Amendment A16. VUS in the ClinVar release of 23 September 2026 (MANE Select missense, disease gene) whose record was reviewed by an expert panel or practice guideline and for which REVEL, BayesDel and AlphaMissense (published ClinGen thresholds) and AVI (Table 1) all gave evidence in the same direction. Comp points, the smallest absolute ACMG/AMP points among the three missense predictors. Scenario, the other evidence with which these points would complete a likely pathogenic (>= +6 points) or likely benign (<= -1 point) classification under the ACMG/AMP point system; it is not a reclassification, because ClinVar does not record the points of the remaining evidence and expert panels may already have applied their own computational criteria. Ev CADD16, CADD v1.7 scores read with the published CADD v1.6 thresholds; Flip AVI, evidence in opposite directions under those thresholds and under calibrated AVI. AF max, gnomAD v4.1 group maximum allele frequency. Ordered by absolute points and genomic position. The 483,690 VUS with concordant evidence at any review status are in the repository (calibration/results/v17_avi/vus_candidates/vus_concordant_evidence.tsv.gz).

*Provided in Supplementary_Tables_S1-S22.xlsx, sheet S16 (1094 rows).*

**Table S17. Calibration of CADD v1.7 and AVI in the intronic splice region.**

Thresholds (one seed, 10,000 bootstrap replicates, missense prior 0.0441) for SNVs 3 to 8 nucleotides into introns of MANE Select transcripts, from 2,392 P/LP and 68,917 B/LB ClinVar variants and 175,221 rare gnomAD variants at the same positions and genes, compared with the missense thresholds.

*Provided in Supplementary_Tables_S1-S22.xlsx, sheet S17 (16 rows).*

**Table S18. Missense-calibrated thresholds in noncoding and synonymous variant classes, and MPRA.**

Reference variants per region class; AUROC of CADD and AVI per class; likelihood ratios (with 1,000-replicate bootstrap intervals) of each missense-calibrated evidence interval in each class and whether the conservative bound meets the required ratio; AUROC for functional versus non-functional SNVs in saturation-mutagenesis MPRA data (Kircher et al. 2019).

*Provided in Supplementary_Tables_S1-S22.xlsx, sheet S18 (170 rows).*

**Table S19. CADD v1.6 versus v1.7 on identical reference data.**

Thresholds (median and range over five seeds) of CADD v1.6 and v1.7 calibrated on the same 2026 reference sets, AUROC with 95% bootstrap intervals and the paired difference, the percentage of reference variants whose evidence level changes between versions, and the full cross-tabulation of evidence levels (each version at its own thresholds).

*Provided in Supplementary_Tables_S1-S22.xlsx, sheet S19 (72 rows).*

**Table S20. Thresholds with and without HGMD Public positions.**

Thresholds of five predictors calibrated on the ClinVar 2019-12-02 and 2026-09-23 intersection sets with all variants (all) and after excluding SNVs at positions of single-nucleotide HGMD Public 2020.4 records (noHGMD), with the published values where available. Set sizes after exclusion were r2019_noHGMD 5395 P/LP, 9681 B/LB, 1665924 gnomAD; r2026_noHGMD 26254 P/LP, 72077 B/LB, 1872367 gnomAD.

*Provided in Supplementary_Tables_S1-S22.xlsx, sheet S20 (40 rows).*

**Table S21. Thresholds across ClinVar releases and before or after November 2022.**

Thresholds (seed 20260925) of five predictors calibrated on the intersection sets of ClinVar releases 2019-12-02, 2022-11-19, 2024-01-27 and 2026-09-23, and on 2026 variants already present in November 2022 (pre2022) or added later (post2022). The common-scale block expresses each threshold as the percentage of 2026 gnomAD intersection variants meeting it (post hoc). Set sizes (P/LP, B/LB) were r2019 17330 / 11379; r2022 32868 / 32044; r2024 40282 / 49090; r2026 59034 / 75306; post2022 30007 / 51383; pre2022 29027 / 23923.

*Provided in Supplementary_Tables_S1-S22.xlsx, sheet S21 (70 rows).*

**Table S22. Calibration by gene class.**

Class sizes, thresholds with 2.5th-97.5th percentile bootstrap intervals, pairwise comparisons (classes considered different when intervals do not overlap) and the percentage of reference variants whose evidence level changes with class-specific instead of global thresholds.

*Provided in Supplementary_Tables_S1-S22.xlsx, sheet S22 (458 rows).*
